# An automated and fast sample preparation workflow for laser microdissection guided ultrasensitive proteomics

**DOI:** 10.1101/2023.11.29.569257

**Authors:** Anuar Makhmut, Di Qin, David Hartlmayr, Anjali Seth, Fabian Coscia

## Abstract

Spatial tissue proteomics integrating whole-slide imaging, laser microdissection and ultrasensitive mass spectrometry is a powerful approach to link cellular phenotypes to functional proteome states in (patho)physiology. To be applicable to large patient cohorts and low sample input amounts, including single-cell applications, loss-minimized and streamlined end-to-end workflows are key. We here introduce an automated sample preparation protocol for laser microdissected samples utilizing the cellenONE® robotic system, which has the capacity to process 192 samples in three hours. Following laser microdissection collection directly into the proteoCHIP LF 48 or EVO 96 chip, our optimized protocol facilitates lysis, formalin de-crosslinking and tryptic digest of low-input archival tissue samples. The seamless integration with the Evosep ONE LC system by centrifugation allows ‘on-the-fly’ sample clean-up, particularly pertinent for laser microdissection workflows. We validate our method in human tonsil archival tissue, where we profile proteomes of spatially-defined B-cell, T-cell and epithelial microregions of 4,000 µm^2^ to a depth of ∼2,000 proteins and with high cell type specificity. We finally provide detailed equipment templates and experimental guidelines for broad accessibility.

## INTRODUCTION

Spatial omics approaches are transforming biomedical research and are gaining increasing momentum for diverse biomedical applications (1, 2). While current spatial profiling methods are dominated by powerful spatial transcriptomics and imaging-based spatial proteomics (SP) concepts (3, 4), SP approaches leveraging the sensitivity and quantitative power of state-of-the-art mass spectrometry (MS)-based proteomics workflows, have been developed more recently (5, 6). Our group co-developed Deep Visual Proteomics (7), which combines subcellular resolution imaging, semi-automated laser microdissection (LMD) and single-cell sensitivity mass spectrometry. Follow-up work based on further improved sample preparation and LC-MS workflows recently led to the first single-cell (i.e. slice of a single cell) proteome measurements from frozen and formalin-fixed (FFPE) tissues (8, 9). In FFPE tissue, we reproducibly quantified up to 2,000 proteins from single excised hepatocyte contours (∼5,000 µm^3^ in volume) based on an optimized sample preparation protocol and label-free based diaPASEF acquisition (10) on the Bruker timsTOF SCP instrument (11). The steadily improving sensitivity of modern LC-MS setups (12), which are key to tissue proteomics applications with further increased spatial resolution, also necessitate the development of robust and automated sample preparation workflows to cope with the growing demand for higher sample throughput, while not compromising sensitivity.

Sample preparation workflows for ultra-low input and single-cell proteomics (SCP) applications rely on automated and loss-reduced robotic workflows. So far, these methods were mostly optimized for cell suspensions combined with cell sorting techniques (e.g. FACS) (13, 14). Here, the cellenONE system was introduced as a versatile platform for label-free (15) or label-based (16–18) SCP capable of nanoliter liquid dispensing and imaging-based single cell sorting in combination with precise temperature and humidity-level controls. Together with a dedicated line of Teflon-based chips containing nano-wells for reduced surface adsorption (i.e. ‘proteoCHIPs’), proteome coverage from trace sample amounts could be significantly improved.

More recently, this workflow has been integrated with the Evosep ONE system based on a new chip design (proteoCHIP EVO 96), thereby entirely omitting additional pipetting steps for sample transfer prior LC-MS acquisition. While this pipeline benefits from low nanoflow gradients for improved sensitivity (e.g. Whisper 40 SPD), it also provides options for higher throughput applications (e.g. 60 SPD or Whisper 80 SPD). For these reasons, we hypothesized that the cellenONE system should also greatly benefit the processing of ultra-low input tissue samples obtained by LMD. However, due to different analytical requirements for sample collection, processing and clean-up of LMD based tissue samples compared to cell suspensions, novel sample preparation protocols are crucial. With this in mind, we here introduce and benchmark an automated sample preparation workflow for ultra-low input laser microdissected (tissue) samples based on the cellenONE system.

## RESULTS

### Integrating laser microdissection with the cellenONE robotic system for spatial proteomics

To enable automated sample preparation of laser microdissected samples with the cellenONE robotic system, we first designed LMD collection plate adapters for the proteoCHIP LF 48 (LF 48 chip) and proteoCHIP EVO 96 (EVO 96 chip). Collection devices were designed for the Leica LMD7 microscope, a state-of-the-art microdissection system that collects samples through gravity. As the collection plate adapter is located between the light source (below collector) and objective (above collector), the use of transparent material such as acryl glass was important to transmit enough light for regular microscopic sample inspection. Our adapter design supports the collection of 144 (three LF 48 chips) and 96 (one EVO 96 chip) samples per LMD session (**Figure 1A, C**). We designed these adapters with minimal distance between the collection device and sample holder to improve tissue collection efficiency. Adapter design and 3D printer templates (.stl file format) are provided in the **Supplementary Figure 1A** and **Supplementary Materials**.

**Figure 1.**
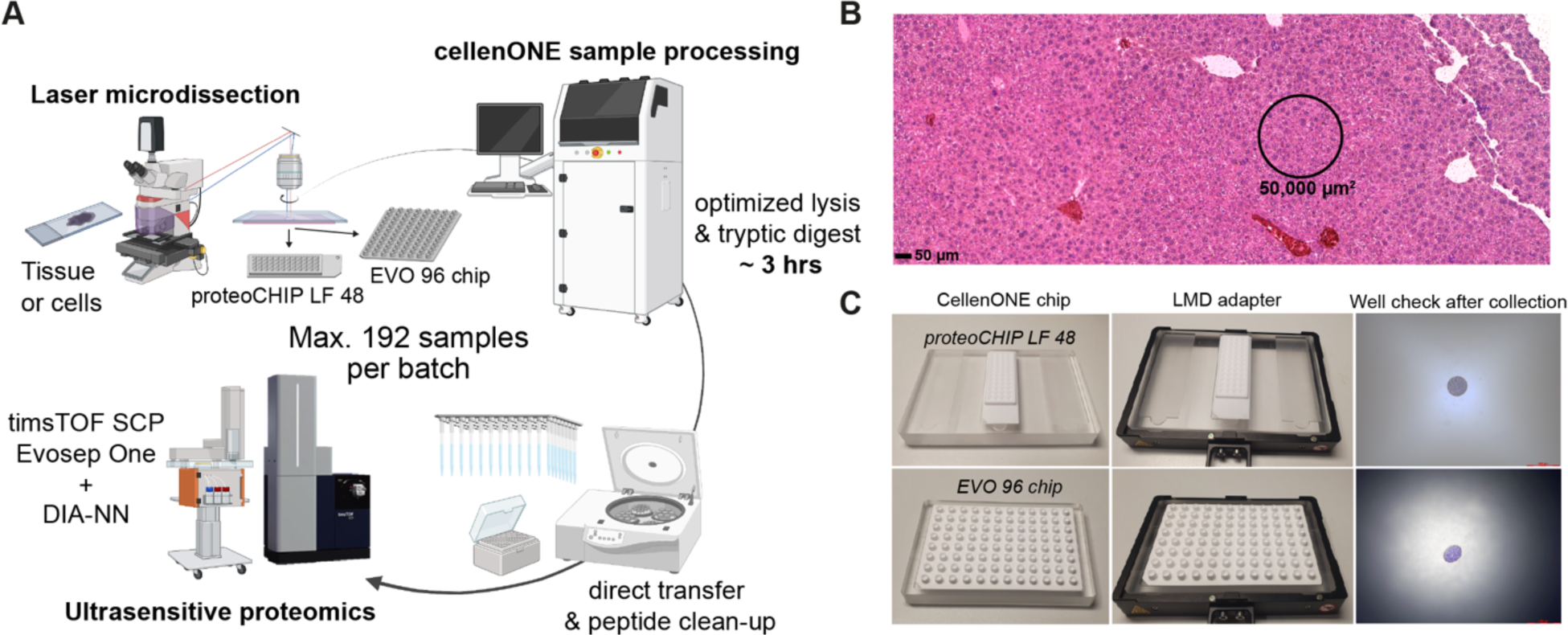
Combining laser microdissection with the cellenONE robotic system for spatial proteomics. **(A)** Overview of the spatial tissue proteomics workflow combining LMD, cellenONE based sample preparation and ultrasensitive LC-MS analysis. **(B)** H&E-stained mouse liver tissue section. A representative contour of 50,000 µm^2^ used for the protocol optimization experiments is shown. Scale bar = 50 µm. **(C)** Overview of the LMD collection plate adapters and tissue inspection after collection into the proteoCHIP LF 48 and EVO 96 chip. Scale bar for well inspection = 400 µm.

We first tested the sample collection directly into the microwells of the two cellenONE chips and used H&E-stained mouse liver tissue to visualize excised samples in brightfield mode before and after cutting. For both chip types, small regions of 50,000 µm^2^ (∼50 hepatocytes, **Figure 1B**) could be easily spotted with the cap-check function of the Leica software without any additional hardware adjustments (**Figure 1C**). Note, for other collection devices such as 96-well or 384-well plates, the default travel distance of the 5x objective does not allow to inspect the well bottom. The narrow well design of the cellenONE chips is thus beneficial to facilitate streamlined well monitoring after sample collection. We conclude that our newly designed adapters are fully compatible with LMD using the LMD7 microscope and potentially other LMD systems based on adjusted adapter layouts. We therefore next proceeded with the testing of different lysis buffer conditions for low-input tissue proteomics.

### Optimizing sample preparation conditions for rapid and ultrasensitive tissue proteomics

Having established a suitable plate holder adapter that integrates the LMD7 with the cellenONE robotic system, we next sought to determine the optimal sample preparation conditions to enable robust and sensitive tissue proteome profiling. A standard label-free single-cell protocol on the cellenONE system encompasses a one-step master mix addition of 100-1000 nL of 0.2% n-dodecyl β-D-maltoside (DDM), 10 ng/uL trypsin and 100 mM TEAB pH 8.5, combined with 2h heating at 50°C. Additionally, reagents and single cells are dispensed in wells pre-filled with oil (hexadecane) to overcome sample evaporation (18). Plate-based oil-free SCP protocols have also been developed based on continuous re-hydration during incubation at elevated temperatures (15). Working with laser microdissected samples mounted on hydrophobic membranes (e.g. polyethylene naphthalate, PEN) prevent the use of oil-prefilled wells. These hydrophobic samples don’t mix well with the aqueous buffer phase, thereby repelling the tissue to the hexadecane/buffer interphase. As an alternative, we therefore incorporated an automated re-hydration strategy to prevent sample evaporation by continuously adding water to each well in a user-definable manner.

As FFPE tissue proteomics workflows generally include prolonged heating to aid formalin de-crosslinking (19–21), we increased the incubation temperature to the instrument’s maximum of 65°C. To evaluate our proteomics results, we then used the same tissue type, sampling amount and LC-MS settings as employed in our recent study (9). We therefore expected that mouse liver tissue contours of 50,000 µm^2^ would result in approx. 4,000 protein groups per sample with high quantitative reproducibility (i.e. Pearson r = 0.95 – 0.99) based on a 15-min active nanoflow gradient, optimal window design diaPASEF method and DIA-NN (22) analysis with a tissue-refined spectral library (**Methods**). For our initial test run, we processed all samples in the proteoCHIP LF 48 and after lysis and digestion, peptides were transferred manually into Evotips for clean-up. Using a one-step protocol (master mix including trypsin), the proteome depth was low and inconsistent, suggesting that the temperature increase from 50°C to 65°C severely affected trypsin activity (**Figure 2A**). This was supported by a doubling of tryptic miscleavage to 50 % and higher (**Supplementary Figure 2A**). We therefore designed an adjusted ‘two-step’ method based on the following ideas: 1) Heating in 0.2% DDM for one hour at 65°C improves lysis and formalin de-crosslinking, 2) separate enzyme addition and digestion carried out at a lowered temperature of 37°C improves digestion, and 3) a continuous re-hydration throughout the method prevents sample evaporation. Incorporating these adaptations, we then identified 25,000 precursors and 3,700 unique proteins per liver tissue microregion, on par with our previous data. This modified method provided an excellent basis for further protocol improvements. We aimed for high protocol robustness, short (2-3 hours) total sample preparation time and a seamless integration with downstream LC-MS. To this end, we designed seven different protocols, which varied in either lysis buffer composition (0.2% DDM, DMSO, or combined), heating duration (1-3 hours) or length of trypsin/LysC digest (1-2 hours) (**Figure 2B, Supplementary Figure 2B**). The low-vapor-pressure solvent DMSO in the lysis buffer was included as an alternative to the continuous re-hydration strategy.

**Figure 2.**
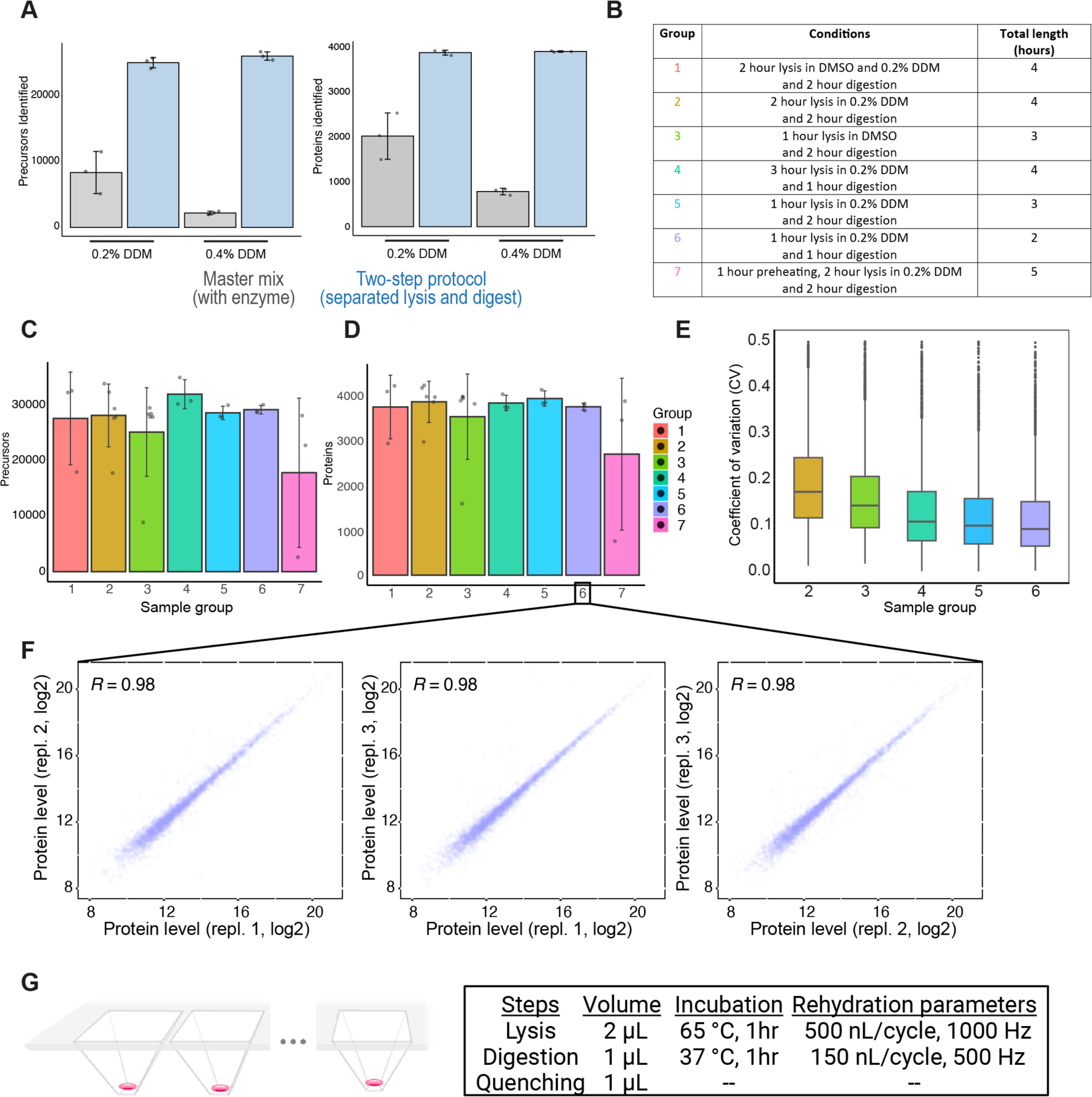
Optimizing sample preparation conditions for rapid and ultrasensitive tissue proteomics. **(A)** Average number of identified precursors (left) and proteins (right) from murine liver tissue samples processed with the standard one-step (grey, master mix including trypsin) or a two-step protocol with separate trypsin addition (blue). Areas of 50,000 µm^2^ were used and two DDM concentrations (0.2% and 0.4%) were compared. Averages are shown from triplicate measurements with standard deviations as error bars. **(B)** Overview of the tested sample preparation conditions. **(C) and (D)** Average number of identified precursors (C) and proteins (D) from diaPASEF measurements of murine liver tissue samples processed with different sample preparation methods. Areas of 50,000 µm^2^ were used. Averages are shown from triplicate measurements for conditions 1, 4, 5, 6 and 7 and six replicates for conditions 2 and 3 with standard deviations as error bars. **(E)** Box plots showing the coefficient of variation (CVs) of protein quantification across different sample preparation methods. CVs were calculated from triplicates (conditions 1, 4, 5 and 6) and six replicates (conditions 2 and 3) of non-log-transformed data. Box plots define the range of the data (whiskers), 25th and 75th percentiles (box), and medians (solid line). **(F)** Proteome correlations (Pearson r) of tissue replicates obtained from sample preparation condition 6. **(G)** Summary of the optimized sample preparation protocol (condition 6), highlighting main parameters including buffer volume, incubation time and re-hydration steps.

Overall, our results revealed more variability with the DMSO based protocols (**Figure 2C-D**) and a somewhat similar number of identified precursors and proteins for the DDM based methods with stepwise lysis and digestion (conditions 2-6). Methods 4 - 6 showed the lowest median protein CVs (11%, 10% and 9% respectively) and a high overlap (>95%) of identified proteins (**Supplementary Figure 2C**). As protocol #6 had the shortest total sample preparation time (2 hours) and lowest median protein CV (9%) at a similar proteome coverage, we chose this condition (1h 65°C lysis in 0.2% DDM + 1h 37°C trypsin/LysC digestion) as our preferred protocol (**Figure 2E-G**).

### Spatially-resolved proteomics of human tonsil tissue

To explore the capacities of our optimized tissue proteomics workflow integrating laser microdissection with the cellenONE system, we performed a proof-of-concept experiment using human tonsil tissue, which is a secondary lymphoid organ and comprised of distinct microanatomical compartments fulfilling diverse adaptive immune-cell functions (23). This tissue type is thus ideal to benchmark our spatial tissue proteomics workflow. In addition, the goal for this experiment was to integrate the EVO 96 chip as this design facilitates the centrifugation-based transfer of peptide samples into Evotips for streamlined sample clean-up and LC-MS injection.

We immunofluorescently stained a 10 µm-thick tissue section for CD19 (B-cells), CD3 (T-cells), panCK (epithelium) and DAPI (DNA) and selected small cell type specific regions of 4,000 µm^2^ for LMD collection into the EVO 96 chip (**Figure 3A**). Following LMD and sample preparation based on the optimized ‘two-hours protocol’ (protocol #6, **Figure 2G**), samples were measured with the Whisper 40 SPD Evosep gradient combined with an optimized diaPASEF method (**Methods**). The entire workflow from lysis to MS-ready Evotips was performed in approx. three hours, drastically reducing sample preparation time compared to previous low-input tissue proteomics workflows (6, 24), including our own (9). Using DIA-NN, we quantified up to 2,000 proteins per sample (**Figure 3B**) with low intra-group protein CVs (**Figure 3C**), which clearly grouped proteomes by cell type (**Figure 3D**). Our data included many known immune cell and functional markers (e.g. STAT1, STAT5A, PARP1, PCNA) and cell type specific cell surface receptors (e.g. CD3D and CD5), which were significantly regulated across the three sample groups (**Figure 3E-F**). Pathway enrichment analysis comparing the B-cell and T-cell proteomes showed a strong enrichment for immunoglobulin and B-cell-mediated immune functions up-regulated in B-cell samples, whereas T-cell regions were characterized by high proteins levels related to the T-cell receptor complex, RAGE receptor, as well several chromatin related terms (**Figure 3G**).

**Figure 3.**
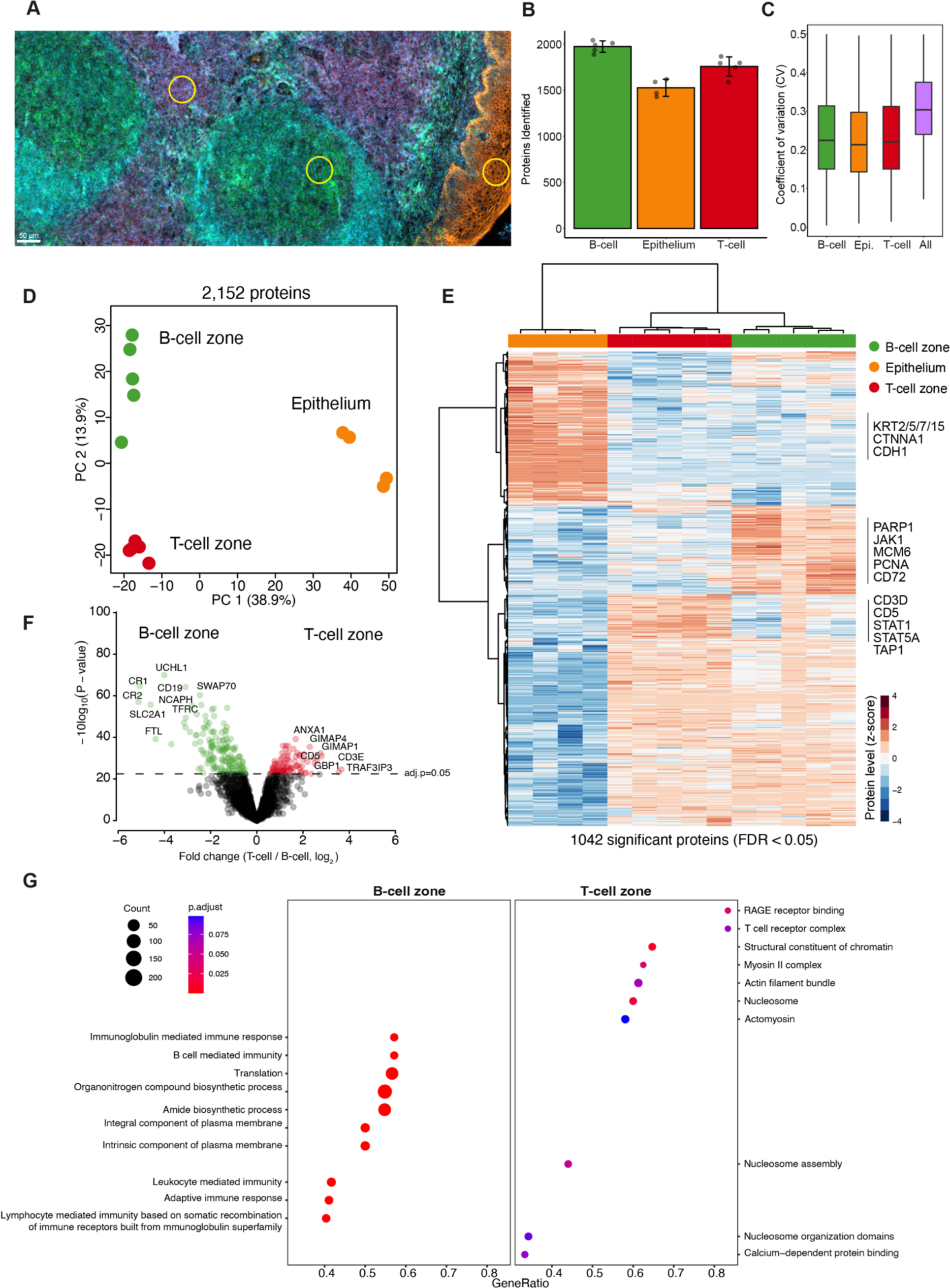
Spatially resolved proteomics of human tonsil tissue. **(A)** Immunofluorescence whole-slide image of a 10-µm thick tonsil tissue section stained for CD3 (T-cells), CD19 (B-cells), pan-CK (epithelium) and DNA (DAPI). Scale bar = 50 µm. Yellow circles illustrate representative contours of 4,000 µm^3^ used for proteomic profiling **(B)** Average number of identified proteins (global protein FDR < 0.05) from each tissue region. Averages are shown from four to five replicates of B-cell, T-cell and epithelial zone samples. **(C)** Coefficient of variation (CV) of protein quantification comparing intra-group versus inter-group variability. **(D)** Principal component analysis of B-cell, T-cell and epithelial zone proteomes based on 2,152 protein groups after data filtering (70% valid values in at least one group). **(E)** Unsupervised hierarchical clustering of 1,042 ANOVA significant proteins (permutation-based FDR < 0.05) from B-cell, T-cell and epithelial zone samples. Heatmap shows relative protein levels (z-score) of up-regulated (red) and down-regulated proteins (blue). **(F)** Volcano plot of the pairwise proteomic comparison between the B-cell and T-cell zones. Significantly regulated proteins are highlighted in green (B-cell zone) and red (T-cell zone). A moderated two-sided t-test was applied, with adjusted p-value of < 0.05. **(G)** Pathway enrichment analysis (Reactome) based on the t-test difference between B-cell and T-cell zone samples using the ClusterProfiler R package. Selected pathways with a Benjamin-Hochberg FDR < 0.1 are shown.

In conclusion, these data demonstrate how our optimized LMD-cellenONE workflow can be applied to gain detailed insights into spatially and cell type resolved-proteomes from minute amounts of archival tissue. Furthermore, our work builds an important framework for future spatial tissue proteomics applications on the basis of streamlined sample processing and ultrasensitive LC-MS.

## DISCUSSION

In this work, we explored the possibility to integrate the Leica LMD7 microscope with the cellenONE robotic system for automated sample processing of low-input laser microdissected samples. To this end, we first designed open-source collection plate adapters for the two commercially available label-free chips (proteoCHIP LF 48 and EVO 96) and successfully evaluated their utility as new tissue collection devices. Our initial tests revealed that current ‘one-step’ protocols (combined buffer for lysis and digestion) developed for single-cell proteomics are suboptimal for FFPE tissue analysis, due to the higher analytical demands to efficiently process crosslinked tissue samples. We therefore undertook a number of adaptations based on our previous results to optimize ultra-low input tissue proteomics workflows (20). We increased the heating temperature to 65°C for enhanced lysis and formalin de-crosslinking, uncoupled lysis and enzymatic digestion to improve tryptic digestion, and added a continuous re-hydration step to prevent sample evaporation in the absence of the hexadecane oil layer. These modifications allowed us to quantify nearly 4,000 proteins from regions of 50,000 µm^2^ mouse liver tissue (50-cell equivalents) with excellent quantitative reproducibility (Pearson r = 0.98) and on par with our previous data using a less automated overnight protocol (9).

We finally applied this optimized workflow to human tonsil FFPE tissue and used the EVO 96 chip for sample preparation. This chip enables streamlined sample clean-up in Evotips after centrifugation-based sample transfer. As sample clean-up steps are particularly important for laser microdissected samples, for example due to left-over membrane pieces that could compromise chromatography performance over time, the EVO 96 chip perfectly combines efficient sample processing with near lossless sample clean-up steps prior LC-MS analysis. In addition, the relatively flat chip design compared to standard 96 or 384-well plates enables a more streamlined well inspection prior proteomics sample preparation. We illustrate the applicability of this setup when profiling B-cell, T-cell and epithelial cell regions of human tonsil tissue, resulting in cell type-specific proteomes that included many known cell surface receptors, immune cell regulators and functional markers.

In its current form, our workflow has the capacity to process 192 (two EVO 96 plates) or even 288 (six LF 48 chips) samples per batch in approx. three hours, from tissue lysis to MS-ready Evotips. Thus, this pipeline strongly improves sample preparation throughput compared to current state-of-the-art low-input tissue workflows, which typically include overnight incubation steps (6, 7, 24). Our work also provides an important framework for future protocol extensions, for example to integrate label-based DIA strategies (25, 26) for further increased MS throughput and proteome coverage. Coupled to Deep Visual Proteomics and other spatial proteomics approaches, we believe that our workflow could pave the way for higher throughput applications, where possibly thousand samples or more could be processed per user on a single day.

In summary, we here describe a robust and automated sample preparation workflow for laser microdissected samples based on the cellenONE robotic system.

## ACKNOWLEDGEMENTS

We would like to thank our colleagues at the Max Delbrück Center (MDC) for their support and fruitful discussions. In particular, we thank Steffen Tornow from the TFM/scientific workshop for his support to design the LMD adapters. Ulrike Stein we thank for her support to perform the mouse liver experiments. Christian Sommer we thank for mass spectrometry support and Simon Schallenberg (Charité Pathology, Berlin) for his help with the tonsil tissue experiments. Furthermore, we acknowledge the MDC technology platform ‘Proteomics’ for their great support. A.M., D.Q. and F.C. acknowledge support by the Federal Ministry of Education and Research (BMBF), as part of the National Research Initiatives for Mass Spectrometry in Systems Medicine, under grant agreement no. 161L0222.

## AUTHOR CONTRIBUTIONS

Conceptualization: all authors.; Methodology: all authors; Experiments: all authors; Data analysis: A.M., D.Q. and F.C.; Figures: A.M., D.Q. and F.C; Supervision: A.S and F.C; Funding acquisition: F.C; Writing the original draft: F.C. All the authors reviewed and edited the manuscript.

## MATERIALS & METHODS

### Mouse liver tissue and H&E staining

The animal experiments were performed in accordance with the United Kingdom Coordinated Committee on Cancer Research (UKCCR) guidelines and were approved by local governmental authorities (Landesamt für Gesundheit und Soziales Berlin, Germany).

The experiments for optimizing sample preparation conditions were done with formalin-fixed paraffin-embedded (FFPE) mouse liver tissue. 6-8 weeks old female C57BL/6 mice from Jackson Laboratory were used. C57BL/6 mice were housed in individually ventilated cages in a specific pathogen-free mouse facility at the Max-Delbrück Center for Molecular Medicine (Berlin, Germany). For liver excision, anesthetized mice were sacrificed by cervical dislocation, and the livers were removed, rinsed twice in ice-cold PBS, and transferred to 4% formaldehyde solution for fixation (fixation for at least 24h to 48h). Thereafter, livers were paraffin-embedded for further histological analyses. The FFPE block was sectioned at 5 µm thickness on PPS frame slides (Leica, 11600294), and left in the oven overnight at 37°C. Before deparaffinization, slides were heated at 60 °C for 10 minutes for better tissue adhesion. Mouse liver samples were stained with hematoxylin-eosin.

### Human tonsil tissue and immunofluorescence staining

The study was performed according to the ethical principles for medical research of the Declaration of Helsinki and approval was obtained from the Ethics Committee of the Charité University Medical Department in Berlin (EA1/222/21).

Human tonsil tissue FFPE blocks were provided by the Institute of Pathology at Charité University Hospital, Campus Mitte. Tonsil tissues were sectioned at 10 µm thickness on PPS frame slides. After tissue adhesion, the samples were subjected to heat-induced (HIER) epitope retrieval to enhance the antibody binding in the further immunofluorescence staining step. Briefly, the samples were heated at 95°C for 20 min and cooled down at room temperature for 30 min. Three conjugated antibodies targeting CD20 (dilution 1:50, Thermofisher, 53-0202-80, Alexa Fluor 488), CD3 (dilution 1:100, Abcam, ab198937, Alexa Fluor 647), pan-cytokeratin (dilution 1:100, Thermofisher, 41-9003-80, eFluor 570) were used to stain the tonsil tissue at 4 °C overnight in a humidity chamber. All antibodies were diluted in Odyssey Blocking Buffer (LI-COR BioScience, 927-70001). Tissues were washed four times in PBS after the antibody incubation, and a subsequent 10 min Hoechst (1:1000 in PBS, Thermo Fisher Scientific, 62249) staining was used for nuclear staining.

Finally, serial ethanol steps (70%, 80%, and 99%) were used to dehydrate the slides and a coverslip was mounted with Diamond anti-fade mounting medium (Invitrogen, cat.no. P36961).

### Design of Leica LMD7 collection plate adapters

Adapters for the proteoCHIP LF 48 and EVO 96 were produced from transparent acryl glass (PMMA-gs) using a CNC milling machine. Production is also possible through conventional 3D printers based on the provided files (.stl format). Design and dimensions are shown in **Supplementary Figure 1A** and 3D printer files provided in the **Supplementary Materials**.

### Whole-slide imaging and laser microdissection

Images of immunofluorescence-labelled tonsil tissue sections were acquired using an Axioscan 7 system (Zeiss), equipped with wide-field optics, a Plan-A photochromat 10x/0.45 M27 objective and a quadruple-band filter set for Alexa fluorescent dyes. The acquired images were imported to QuPath (version 0.4.3) for further annotation. The different regions of interest were annotated in the software and exported in a geojson format together with three reference points for contour alignment.

The annotations in a geojson format were translated to the required .xml format for laser microdissection. The contours were assigned to different target locations on proteoCHIP LF 48 and proteoCHIP EVO 96. The code for processing the shapes is available at github.com/CosciaLab/Qupath_to_LMD, it uses geopandas and the py-lmd package (27).

Laser microdissection was done with the Leica LMD7 system (Leica Laser Microdissection software V 8.3.0.08259). 3D printed plate holders were used to adjust the location and height of proteoCHIPs LF 48 and EVO 96 in the laser microdissection system and customized plate layouts were defined by using the universal holder function in the LMD software. Depending on the contour size, tissue was cut with a 5x, 20x or 40x objective in fluorescence or brightfield mode.

### Sample preparation using the cellenONE system

Contours were cut and sorted into proteoCHIP LF 48 and proteoCHIP EVO 96. To concentrate tissue samples at the bottom of the LF 48 and EVO 96 chips, 10 µL of acetonitrile was added to each well after collection and vacuum dried (15 min at 60°C). Another well inspection is recommended before proteomics sample preparation to ensure high collection efficiency.

For the preparation of all reagents and buffers, purified and filtered water (>18 MΩ, <3ppb TOC at 25 °C) was used. 2 µl of lysis buffer (0.2% DDM, 0.1M TEAB pH 8.5) was dispensed into each well using the pump function of the cellenONE (Cellenion, France) at 8°C and 50% humidity. After dispensing, the temperature was increased to 65°C and humidity to 85%. Continuous re-hydration (500 nL/cycle, 1000Hz) was activated to prevent evaporation of lysis buffer from the wells. The volume dispensed per cycle might need to be adjusted depending on the local temperature and humidity level to avoid complete evaporation. After 60 min incubation, the temperature was set to 20°C, and re-hydration was continued until the temperature of 25°C was reached. At 20°C, the run was stopped. 1 µL of enzyme mix (lysC & trypsin, 10 ng/µL in water, Promega, Cat. V5072) was dispensed into each well by using the pL-volume dispensing function of the cellenONE at 20°C and 85% humidity. After dispensing, the temperature was increased to 37°C and humidity remained at 85%. The reaction mixture was incubated for 1 hour with continuous re-hydration (150 nL/cycle, 500 Hz), after which digestion was stopped by acidification using 2.5 µL 0.1% trifluoroacetic acid (TFA) per well.

### Peptide clean-up with C-18 tips

After digestion, Evotip (Evosep, Odense, Denmark)-based peptide clean-up was performed as recommended by the manufacturer. Briefly, 20 µL of buffer B (99.9% ACN, 0.1% FA) was added to each C-18 tip (EV2013, Evotip Pure, Evosep) and centrifuged at 700 rcf for 1 min. Then, 20 µL of buffer A (99.9% water, 0.1% FA) was added from the top of each C-18 tip, activated in isopropanol for 20 seconds and centrifuged again at 700 rcf for 1 min. For proteoCHIP LF 48, digested tissue samples were manually loaded from the chip onto Evotips, washed once with 20 µL buffer A and finally eluted with 20 µL buffer B to a 96-well plate (Thermo Fisher Scientific, AB1300), and vacuum dried (15 min at 60°C). Samples were stored at −20°C until LC–MS analysis. Samples produced in the proteoCHIP EVO 96 were transferred directly by mounting the chip on-top of 96 activated Evotips in a Evotip box followed by centrifugation at 700 rcf for 1 min and a washing step with 20 µL buffer A, again at 700 rcf for 1 min. Finally, 100 µL of buffer A was added to each tip and centrifuged at 700 rcf for 10 seconds to move the liquid down to the membrane. Tips then were placed in the tray to a holder filled with buffer A, so that tips are submerged and do not dry.

### Liquid chromatography–mass spectrometry (LC – MS) analysis

For comparison to our recent study (9), initial experiments (Figures 1 and 2) were performed using an EASYnLC-1200 system (Thermo Fisher Scientific) connected to a trapped ion mobility spectrometry quadruple time-of-flight mass spectrometer (timsTOF SCP, Bruker Daltonik). A 15-min active gradient (21 min total) was used with home-packed HPLC columns (20cm x 75um, 1.9-µm ReproSil-Pur C18-AQ silica beads, Dr. Maisch) kept at 40 °C during acquisition. Buffer A consisted of 0.1% formic acid in LC-MS grade water and buffer B is 0.1% formic acid in 90% acetonitrile.

Tonsil tissue proteomes (Figure 3) were acquired on an Evosep ONE (Evosep Biosystems) LC system coupled to the timsTOF SCP. Peptides were separated with Whisper 40 SPD method. A standardized 31-min gradient was run with Aurora Elite column (15cm x 75um, 1.7um, IonOpticks) kept at 50 °C. The mobile phases contain 0.1% FA in water for buffer A and 0.1% FA in acetonitrile for buffer B. For dia-PASEF analysis, we used a dia-PASEF method with 8 TIMS ramps with 3 mass ranges per ramp covering a 400-1000 m/z range by 25 Th windows and an ion mobility range from 0.64 to 1.37 Vs cm^-2^. The mass spectrometer was operated in high sensitivity mode, with an accumulation and ramp time at 100 ms, the collision energy as a linear ramp from 20 eV at 1/K0 = 0.6 Vs cm^-2^ to 59 eV at 1/K0 = 1.6 Vs cm^-2^. The spray voltage in dia-PASEF method was set to 1500 V when running with 21 minutes gradient on EASYnLC-1200 system, and was segmented throughout the gradient with 1500 V at the beginning and end, whereas 1400 V was applied from 5-27 min for Whisper 40 SPD.

### Proteomics data processing and statistical analysis

We used DIA-NN (version 1.8.1) for dia-PASEF raw file analysis and spectral library generation. DIA-NN in silico predicted libraries were generated by providing the human or mouse FASTA file and frequently found contaminants (28) (UP000000589_10090 and UP000005640_9606, downloaded on April 10th and April 8th 2022 respectively). Project-specific libraries were then generated by refining in silico libraries with 20-50 ‘higher-load’ raw files from the same tissue type (mouse liver tissue or human tonsil tissue). The refined murine liver library consisted of 68,006 precursors, 61,554 elution groups and 8,225 protein groups. The refined human tonsil library consisted of 47,999 precursors, 44,675 elution groups and 8,137 protein groups. Raw files were then searched with these refined libraries and using the same DIA-NN version. DIA-NN was operated in the default mode with minor adjustments. Briefly, precursors charge state was set to 2-4, precursor m/z range to 400 – 1000, MS1 and MS2 accuracies to 15.0 ppm, scan windows to 0 (assignment by DIA-NN), isotopologues and MBR were enabled, heuristic protein inference and no shared spectra. Proteins were inferred from genes, neural network classifier was set to single-pass mode, quantification strategy as ‘Robust LC (high precision)’. Cross-run normalization was set to ‘RT-dependent’, library generation as ‘smart profiling’, speed and Ram usage as ‘optimal results’.

Proteomics data analysis was performed with Perseus (29) (version 1.6.15.0) and within the R environment (https://www.r-project.org/, version 4.2.2) with the following packages: *ggplot2 (v3.4.2), FactoMineR (v2.8), factoextra (v 1.0.7.999), reshape2 (1.4.4), viridis (v0.6.3), clusterProfiler (v4.6.2), ReactomePA (v1.42.0), org.Hs.eg.db (v3.16.0).* The Shiny app Protigy (https://github.com/broadinstitute/protigy) was used for data analysis in Figure 3.

For differential expression analysis (t-test or ANOVA), data were filtered to keep only proteins with 70% non-missing data in at least one group. Missing values were then imputed based on a normal distribution (width = 0.3, downshift = 1.8) before statistical testing. For multi-group (ANOVA) or pairwise proteomic comparisons (two sample moderated t-test), a permutation-based and Benjamini-Hochberg based FDR of 5% was applied, respectively, to correct for multiple hypothesis testing. Pathway enrichment analysis (Figure 3G) was performed with the clusterProfiler R package(30). The fold change between B and T-cell samples was used as input variable for the enrichment analysis with a minimum category size of 5 proteins and a maximum of 500. Gene ontology was set to ‘all’ to include terms related to cellular compartments (CC), molecular function (MF) and biological processes (BP). The p-value cut-off was set to 0.1 and pAdjustMethod to “fdr”.

## Data availability

The mass spectrometry proteomics data have been deposited to the ProteomeXchange Consortium (http://proteomecentral.proteomexchange.org) via the PRIDE partner and will be accessible upon publication.

---

**Supplementary Figure 1.**
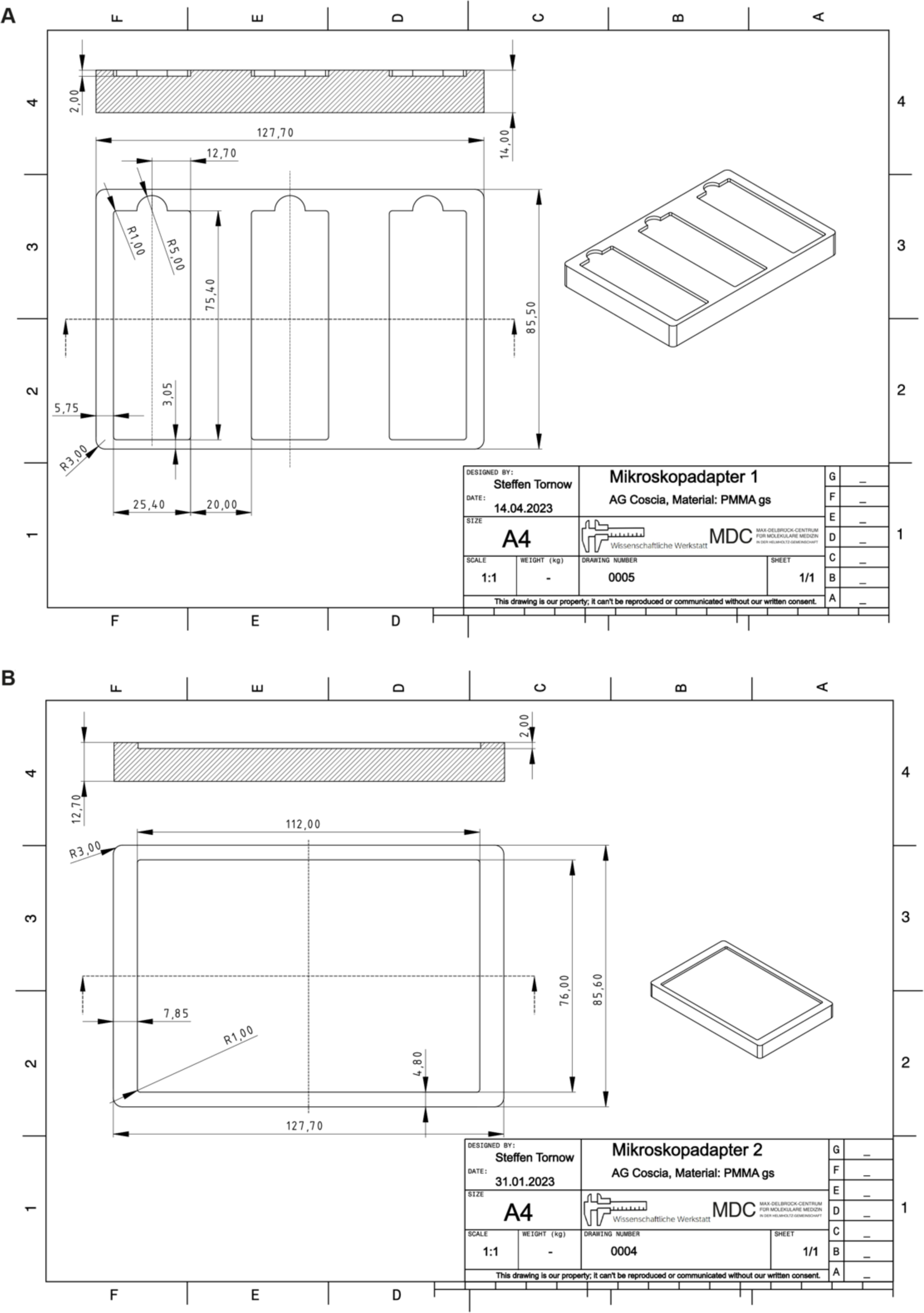
Leica LMD7 collection plate adapter design for the cellenONE proteoCHIPs LF 48 and EVO 96.

**Supplementary Figure 2.**
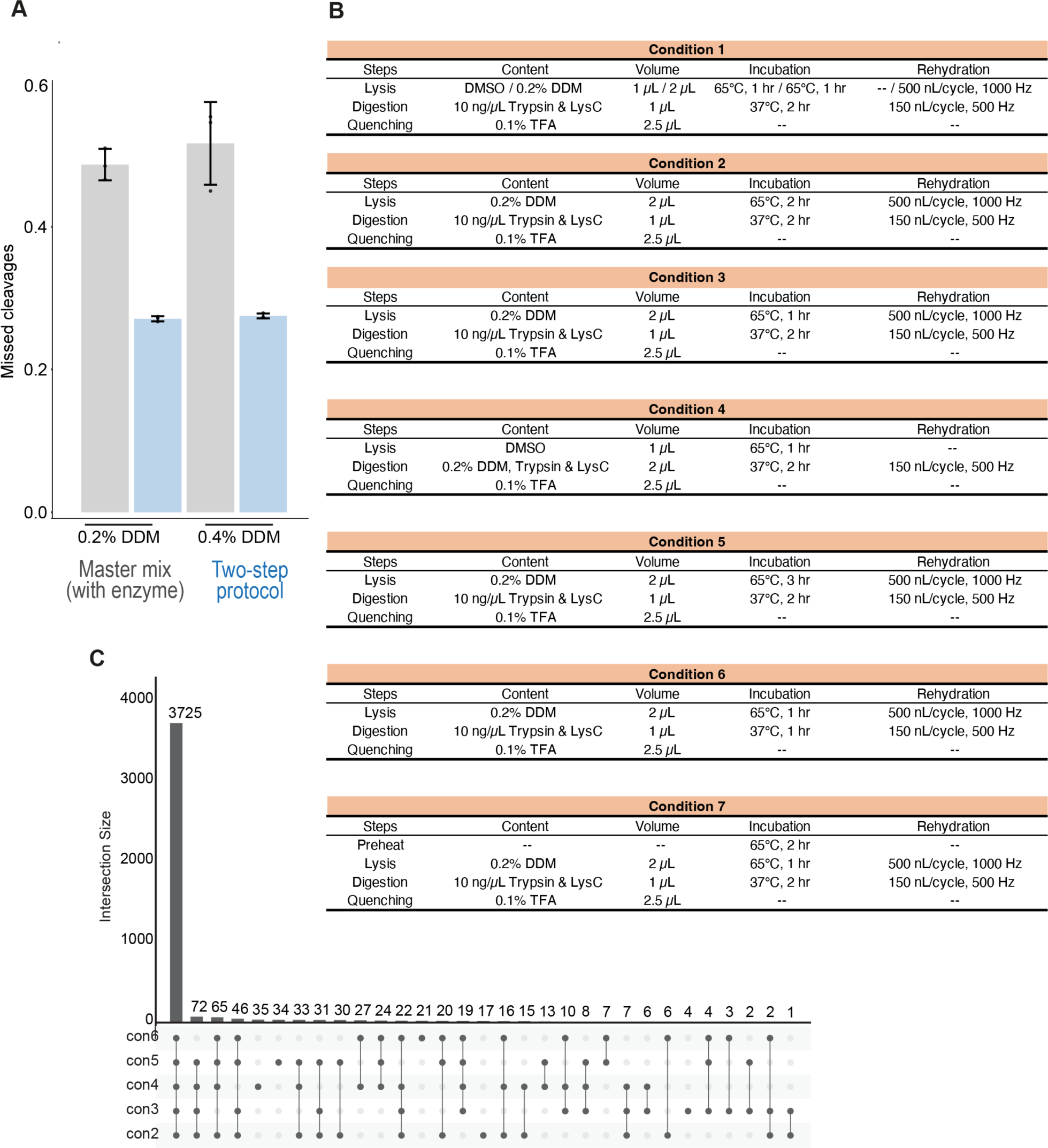
Optimizing sample preparation conditions for tissue proteomics on the cellenONE system. **(A)** Tryptic miscleavage rates of mouse liver tissue samples processed with four different protocols. Grey bars show the results for the one-step protocol with 0.2% and 0.4% DDM and light blue bars based on a two-step protocol (separated lysis and digestion). Data show mean values from triplicates with standard deviations as error bars. **(B)** Overview of seven different sample preparation protocols tested on the cellenONE system. Laser microdissected mouse liver tissue (50,000 µm^2^) was used as test tissue. (C) Upset plot of protein identifications obtained from mouse liver tissue samples processed with the best five protocols (conditions 2-6), depicted in panel (B).

## REFERENCES

1. Vandereyken, K., Sifrim, A., Thienpont, B., and Voet, T. (2023) Methods and applications for single-cell and spatial multi-omics. Nat Rev Genet 24, 494–515

2. Bressan, D., Battistoni, G., and Hannon, G. J. (2023) The dawn of spatial omics. Science (1979) 381, 1–35

3. Lewis, S. M., Asselin-Labat, M. L., Nguyen, Q., Berthelet, J., Tan, X., Wimmer, V. C., Merino, D., Rogers, K. L., and Naik, S. H. (2021) Spatial omics and multiplexed imaging to explore cancer biology. Nat Methods 18, 997–1012

4. Longo, S. K., Guo, M. G., Ji, A. L., and Khavari, P. A. (2021) Integrating single-cell and spatial transcriptomics to elucidate intercellular tissue dynamics. Nat Rev Genet 22, 627–644

5. Mund, A., Brunner, A.-D., and Mann, M. (2022) Unbiased spatial proteomics with single-cell resolution in tissues. Mol Cell 82, 2335–2349

6. Piehowski, P. D., Zhu, Y., Bramer, L. M., Stratton, K. G., Zhao, R., Orton, D. J., Moore, R. J., Yuan, J., Mitchell, H. D., Gao, Y., Webb-Robertson, B. J. M., Dey, S. K., Kelly, R. T., and Burnum-Johnson, K. E. (2020) Automated mass spectrometry imaging of over 2000 proteins from tissue sections at 100-μm spatial resolution. Nat Commun 11,

7. Mund, A., Coscia, F., Kriston, A., Hollandi, R., Kovács, F., Brunner, A.-D., Migh, E., Schweizer, L., Santos, A., Bzorek, M., Naimy, S., Rahbek-Gjerdrum, L. M., Dyring-Andersen, B., Bulkescher, J., Lukas, C., Eckert, M. A., Lengyel, E., Gnann, C., Lundberg, E., Horvath, P., and Mann, M. (2022) Deep Visual Proteomics defines single-cell identity and heterogeneity. Nature Biotechnology 2022, 1–10

8. Rosenberger, F. A., Thielert, M., Strauss, M. T., Schweizer, L., Ammar, C., Mädler, S. C., Metousis, A., Skowronek, P., Wahle, M., Madden, K., Gote-Schniering, J., Semenova, A., Schiller, H. B., Rodriguez, E., Nordmann, T. M., Mund, A., and Mann, M. (2023) Spatial single-cell mass spectrometry defines zonation of the hepatocyte proteome. Nat Methods 20, 1530–1536

9. Makhmut, A., Qin, D., Fritzsche, S., Nimo, J., König, J., and Coscia, F. (2023) A framework for ultra-low-input spatial tissue proteomics. Cell Syst 14, 1002–1014.e5

10. Meier, F., Brunner, A. D., Frank, M., Ha, A., Bludau, I., Voytik, E., Kaspar-Schoenefeld, S., Lubeck, M., Raether, O., Bache, N., Aebersold, R., Collins, B. C., Röst, H. L., and Mann, M. (2020) diaPASEF: parallel accumulation–serial fragmentation combined with data-independent acquisition. Nat Methods 17, 1229–1236

11. Brunner, A., Thielert, M., Vasilopoulou, C., Ammar, C., Coscia, F., Mund, A., Hoerning, O. B., Bache, N., Apalategui, A., Lubeck, M., Richter, S., Fischer, D. S., Raether, O., Park, M. A., Meier, F., Theis, F. J., and Mann, M. (2022) Ultra-high sensitivity mass spectrometry quantifies single-cell proteome changes upon perturbation. Mol Syst Biol 18, 1–15

12. Petrosius, V., and Schoof, E. M. (2023) Recent advances in the field of single-cell proteomics. Transl Oncol 27, 101556

13. Budnik, B., Levy, E., Harmange, G., and Slavov, N. (2018) SCoPE-MS: Mass spectrometry of single mammalian cells quantifies proteome heterogeneity during cell differentiation 06 Biological Sciences 0601 Biochemistry and Cell Biology 06 Biological Sciences 0604 Genetics. Genome Biol 19, 1–12

14. Schoof, E. M., Furtwängler, B., Üresin, N., Rapin, N., Savickas, S., Gentil, C., Lechman, E., Keller, U. auf dem, Dick, J. E., and Porse, B. T. (2021) Quantitative single-cell proteomics as a tool to characterize cellular hierarchies. Nat Commun 12, 3341

15. Matzinger, M., Müller, E., Dürnberger, G., Pichler, P., and Mechtler, K. (2023) Robust and Easy-to-Use One-Pot Workflow for Label-Free Single-Cell Proteomics. Anal Chem 95, 4435–4445

16. Leduc, A., Huffman, R. G., Cantlon, J., Khan, S., and Slavov, N. (2022) Exploring functional protein covariation across single cells using nPOP. Genome Biol 23, 1–31

17. Woo, J., Williams, S. M., Markillie, L. M., Feng, S., Tsai, C. F., Aguilera-Vazquez, V., Sontag, R. L., Moore, R. J., Hu, D., Mehta, H. S., Cantlon-Bruce, J., Liu, T., Adkins, J. N., Smith, R. D., Clair, G. C., Pasa-Tolic, L., and Zhu, Y. (2021) High-throughput and high-efficiency sample preparation for single-cell proteomics using a nested nanowell chip. Nat Commun 12, 1–11

18. Ctortecka, C., Hartlmayr, D., Seth, A., Mendjan, S., Tourniaire, G., Mechtler, K., and Biocenter, V. (2022) An automated workflow for multiplexed single-cell proteomics sample preparation at unprecedented sensitivity. bioRxiv, 2021.04.14.439828

19. Magdeldin, S., and Yamamoto, T. (2012) Toward deciphering proteomes of formalin-fixed paraffin-embedded (FFPE) tissues. Proteomics 12, 1045–1058

20. Coscia, F., Doll, S., Bech, J. M., Schweizer, L., Mund, A., Lengyel, E., Lindebjerg, J., Madsen, G. I., Moreira, J. M. A., and Mann, M. (2020) A streamlined mass spectrometry–based proteomics workflow for large-scale FFPE tissue analysis. Journal of Pathology 251, 100–112

21. Kawashima, Y., Kodera, Y., Singh, A., Matsumoto, M., and Matsumoto, H. (2014) Efficient extraction of proteins from formalin-fixed paraffin-embedded tissues requires higher concentration of tris(hydroxymethyl)aminomethane. Clin Proteomics 11,

22. Demichev, V., Messner, C. B., Vernardis, S. I., Lilley, K. S., and Ralser, M. (2020) DIA-NN: neural networks and interference correction enable deep proteome coverage in high throughput. Nat Methods 17, 41–44

23. De Silva, N. S., and Klein, U. (2015) Dynamics of B cells in germinal centres. Nat Rev Immunol 15, 137–148

24. Davis, S., Scott, C., Ansorge, O., and Fischer, R. (2019) Development of a Sensitive, Scalable Method for Spatial, Cell-Type-Resolved Proteomics of the Human Brain. J Proteome Res 18, 1787–1795

25. Thielert, M., Itang, C., Ammar, C., Schober, F., Bludau, I., Skowronek, P., Wahle, M., Zeng, W.-F., Zhou, X.-X., Brunner, A.-D., Richter, S., Theis, F. J., Steger, M., and Mann, M. (2022) Robust dimethyl-based multiplex-DIA workflow doubles single-cell proteome depth via a reference channel Author list. bioRxiv, 1–51

26. Derks, J., Leduc, A., Huffman, R. G., Specht, H., Ralser, M., Demichev, V., and Slavov, N. (2021) Increasing the throughput of sensitive proteomics by plexDIA.

27. Schmacke, N. A., Mädler, S. C., Wallmann, G., Metousis, A., Bérouti, M., Harz, H., Leonhardt, H., Mann, M., and Hornung, V. (2023) SPARCS, a platform for genome-scale CRISPR screening for spatial cellular phenotypes. bioRxiv, 2023.06.01.542416

28. Frankenfield, A. M., Ni, J., Ahmed, M., and Hao, L. (2022) Protein Contaminants Matter: Building Universal Protein Contaminant Libraries for DDA and DIA Proteomics. J Proteome Res 21, 2104–2113

29. Tyanova, S., Temu, T., Sinitcyn, P., Carlson, A., Hein, M. Y., Geiger, T., Mann, M., and Cox, J. (2016) The Perseus computational platform for comprehensive analysis of (prote)omics data. Nat Methods 13, 731–740

30. Yu, G., Wang, L. G., Han, Y., and He, Q. Y. (2012) ClusterProfiler: An R package for comparing biological themes among gene clusters. OMICS 16, 284–287

